# Histopathological effects and micronucleus assay of glyphosate-based herbicides on cultured african catfish (*Clarias Gariepinus, Burchell 1822*)

**DOI:** 10.1101/2021.08.25.457628

**Authors:** S.A. Alarape, E.O. Adebiyi, O.K. Adeyemo

## Abstract

Glyphosate, a brand of agricultural herbicides which intensive use has led to widespread contamination of different ecosystems. This study was designed to determine both organotoxicity and genotoxicity of glyphosate on African Catfish (*Clarias gariepinus*) exposed to different concentrations for 96 hours. Questionnaires were administered (physically and online) to determine the type of glyphosate-based herbicides mostly used by fish farmers. Seventy-five apparently healthy adult *Clarias gariepinus* (300g) were sourced from a local farmer, transported in a plastic keg to Fish and Wildlife Laboratory at the Department of Veterinary Public Health and Preventive medicine for two (2) weeks of acclimatization. After acclimatization, they were further divided into four (4) groups (T0 (Control), T1 (0.003ml/L), T2 (0.0045ml/L) and T3 (0.006ml/L)) by simple randomization and each group replicated into three (3) treatments. At the expiration of 96 hours of exposure, blood samples and organs (Gills, Kidney, and Liver) were collected for mononuclear assay and histopathological lesions respectively.

Exposed groups showed erratic swimming, splashing, and restlessness. Mortalities rate was dose-dependent (two (2) mortalities at 0.0045ml/L concentration (T2) and five (5) mortalities 0.006ml/L concentration (T3)). Observed histopathological lesions occurred at higher dose treatment (0.0045ml/L (T2) and (0.006ml/L) T3)) concentrations. The gills showed diffuse stunted and eroded secondary lamellae and severe congestion of the blood channel at the core of the primary lamellae. Lesions in the Liver include severe diffuse vacuolation of the hepatocytes, moderate to severe portal congestion and mild diffuse vacuolation of hepatocytes and moderate diffuse vacuolation of hepatocytes, and severe portal congestion. In the kidney, there was mild to moderate congestion of the interstitium and focus of interstitial oedema within the parenchyma. There was presence of micronucleus in the fish nucleated red blood cells at higher dose concentrations.

This study showed that Glyphosate-based herbicides are highly toxic to *Clarias gariepinus*, therefore their use near the fish farm or in areas close to the aquatic environment should be discouraged. The agricultural community should also be conscious of the potentially adverse effects of pesticides. This is to prevent the water body from the residue of herbicides that would have washed down to the water.

## Introduction

According to Decline [1], fish production of a pond is the amount of fish gained at harvest and is dependent on the farmer’s management and system of practice. Aquaculture practice is the adoption of innovations in fish farming in an enclosure. The degree of aquaculture practices affects productivity, income, and subsequent development. By implication, the type of production practice adopted indicates the level of technology input to stimulate production growth and sustainability. The sustainability of aquaculture practice depends on the economic disposition and value it adds to the welfare of the farmer [1].

With the rise in fish farming and aquaculture, there has been an exponential increase in the use of agrochemicals or agrichemical. Agrochemicals refer to pesticides including insecticides, herbicides, and fungicides. Such agrochemicals are marketed as harmless, with little or no side effects by the manufacturers. Despite the rapidly increasing use of these agrochemicals, researchers have viewed some of the chemicals as materials whose environmental fate is poorly understood [3].

The use of agrochemicals, such as herbicides in farming and fish farming, most especially around ponds has raised a lot of questions as regards public health. Examples of chemicals with negative effects on target organisms and public health implications used in aquaculture include Copper sulphate, Malachite green, Formalin, Potassium permanganate, etc. According to Alarape et al., [4], *Clarias gariepinus’* exposure to copper sulphate resulted in necrotic ovaries, matted lamellae of the gills, and multifocal severe degeneration of the seminiferous tubules while those exposed to Malachite green resulted in disrupted and depleted seminiferous tubules, focal localized vacuolation of skin, and generalized fatty degeneration of liver [5]. The use of agrochemicals can easily be washed into ponds and other water bodies by rain, altering the physiological parameters of the water and the chemicals could also build up in the fishes’ systems, and this would pose a threat to human health and public health in general. The herbicide glyphosate (N-phosphonomethyl glycine), is a biocide with a broad-spectrum activity introduced for weed control in agricultural production fields in 1974 [6]. Glyphosate is the active ingredient in the Roundup (Monsanto Company, St. Louis) brand of agricultural herbicides and a variety of other herbicide formulations. These formulations which are used widely in agricultural, forestry, and residential markets provide non-selective, post-emergent control of annual and perennial weeds [7]. Glyphosate is one of the common herbicides used as a non-selective herbicide and for aquatic weed control in fish ponds, lakes, canals, slow running water, which is said to be slightly toxic to mammals and fish [8]. Glyphosate is taken up by the foliage of plants and transported throughout the plant resulting in plant death after several days. Glyphosate is formulated with various adjuvants [9], in particular surfactants such as polyoxyethylene amine (POEA), to enhance the uptake and translocation of the active ingredient in plants. The best-known product formulated with polyoxyethylene amine is Roundup® [6].

Glyphosate is the most extensively used herbicide worldwide. The intensive use of glyphosate has led to its widespread contamination of different ecosystems where it influences plants, microorganisms, animals, and many components of the food chain [10]. Glyphosates are major contaminants of rivers and surface waters [11] as well as organisms, including humans, but also food, feed, and ecosystems [7]. Their use and presence in the food chain are further increased again with more than 75% of genetically modified edible plants that have been designed to tolerate high levels of these compounds [12], commercialized in various formulations [13]. Glyphosate is one of the widely used herbicides that could be persistent and mobile in soil and water, and it is known to be one of the most common terrestrial and aquatic contaminants [14]. Glyphosate alone is not used as an herbicide [15]; in fact, it is always blended with different surfactants to increase its perforation into plant cells which adds to its toxicity [16]. It is used to inhibit weed and clear the space for the growth of vegetation in fields apart from enhancing the plantation in the parks, forests, railway lines, public streets, and gardens.

In the past, glyphosate was considered a nontoxic herbicide because of its low LD50 (the concentration that caused 50% deaths); > 4g/kg [17]. Ayoola [18] reported that glyphosate toxicity increased with increased concentration. And according to Gill *et al*., [19], glyphosate’s toxicity is not only observed in unicellular organisms but also shows its toxic effects on many multicellular organisms found on both soil and water.

Toxicological effects like genotoxicity, cytotoxicity, nuclear aberration, hormonal disruption, chromosomal aberrations, and DNA damage have also been observed in higher vertebrates like humans [19]. According to Neibor and Richardson [20], the level of toxicity of any pesticide depends on its bioaccumulation, the different chemistries of the compound forming the pesticide, and the reactions of the organisms receiving the toxicant.

## Methods

Fifty pre-tested structured questionnaire asides from online Google forms were distributed to fish farmers to determine the usage of glyphosate chemicals on fish farms as well as the glyphosate brand commonly used. Seventy-five apparently healthy adult *Clarias gariepinus* (300g) were sourced from a local farmer. They were transported in a plastic keg to the Fish and Wildlife Laboratory at the Department of Veterinary Public Health and Preventive medicine that same day. The fishes were subsequently transferred into holding tanks for two (2) weeks acclimatization and were fed protein pellets daily at 5% body weight and later divided into four (4) groups by simple randomization. There were four (T_1_, T_2_, T_3_ and T_0_) concentrations (0.15mls (0.003ml/L), 0.225mls (0.0045ml/L) and 0.3mls (0.006ml/L)) and a control of (0ml) respectively and three replicas of each concentration. The chemicals at different concentrations were added to a 50L mark tank of water respectively and aerated approximately one hour daily. Physical observation and behavioural changes were observed after the introduction of Force-Up. Water quality parameters were taken using Farmer’s test kit (for Alkalinity, pH, Nitrite, and Water hardness), Dissolved Oxygen meter (for Dissolved Oxygen, Temperature, Atmospheric Pressure, and Biochemical Oxygen demand), and Photometer (for ammonia concentration). Fresh solutions were prepared every twenty-four hours while aeration and water quality parameters were repeated daily for five days.

### Sample Collection

Blood samples were collected from the caudal veins and poured inside EDTA embedded bottles. Organs (gills, liver, and kidney) were also collected through dissection of the fish and placed in plain labeled bottles for histopathology, and preserved with Bouin’s fluid for proper tissue preparation and slide production.

### Ethical approval

The University of Ibadan Animal Care and Use in Research Committee (UI-ACUREC) approved this study protocol and all procedures with an assigned number UI-ACUREC/019-0220/6.

## Results

Out of fifty questionnaires administered physically, only thirty (30) were retrieved while there were eight (8) respondents from online Google form. The results of the analysis showed that Catfish farmers have the highest number (Fig 1) of respondents (25 (65.8%)), followed by Heterobranchus (7 (18.4%)) and Hetero-Clarias (6 (15.8%)). Force-Up had the highest usage (13 (18.42%)), while Vinash had the least 3 (7.89%) use among the respondents, though 9 (23.68%) of the respondents claimed they were not using herbicides in the control of weeds on the farm. (Fig. 2).

**Fig. 1.**
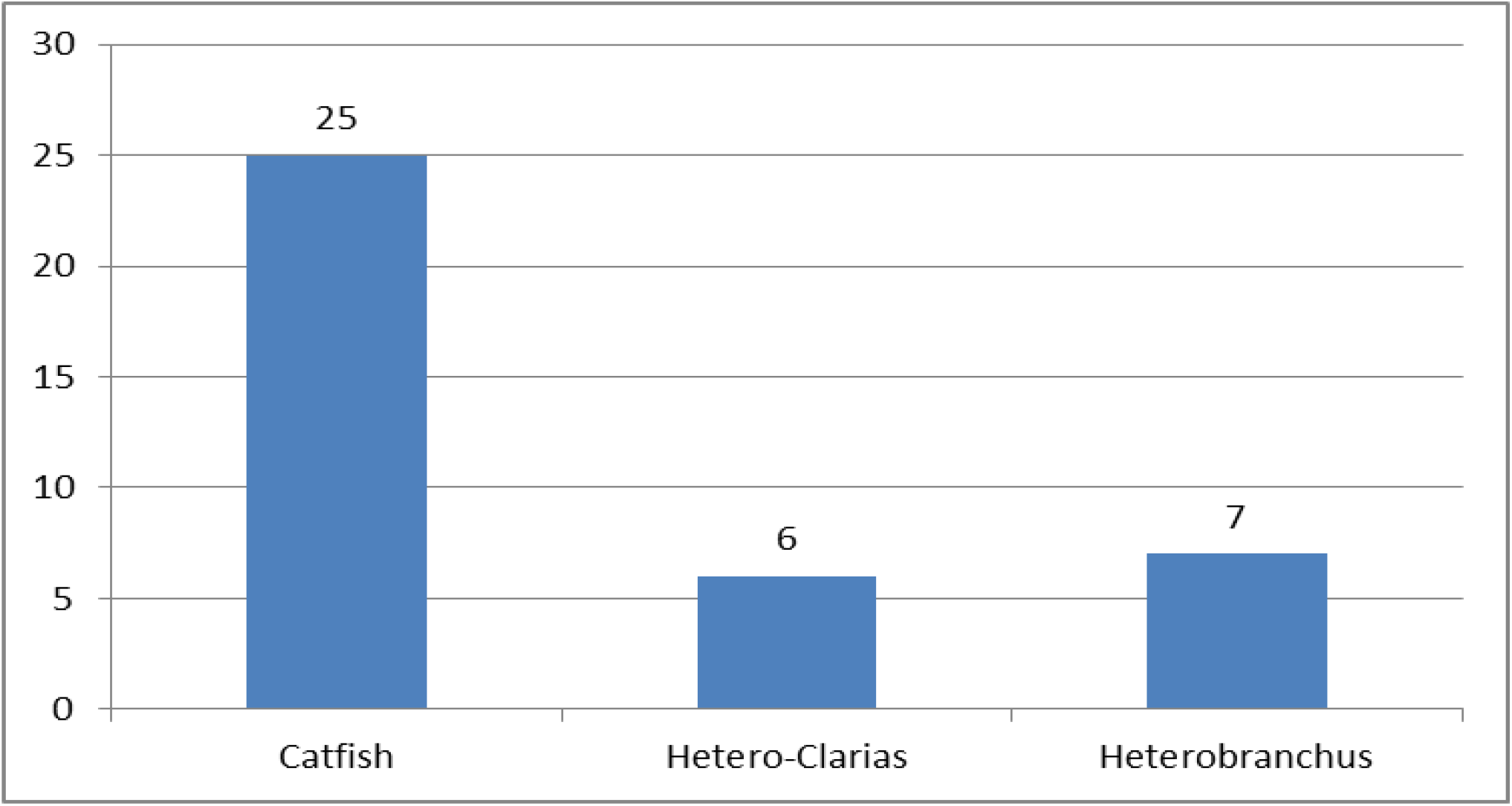
showing Distribution of Cultured Fish.

**Fig. 2.**
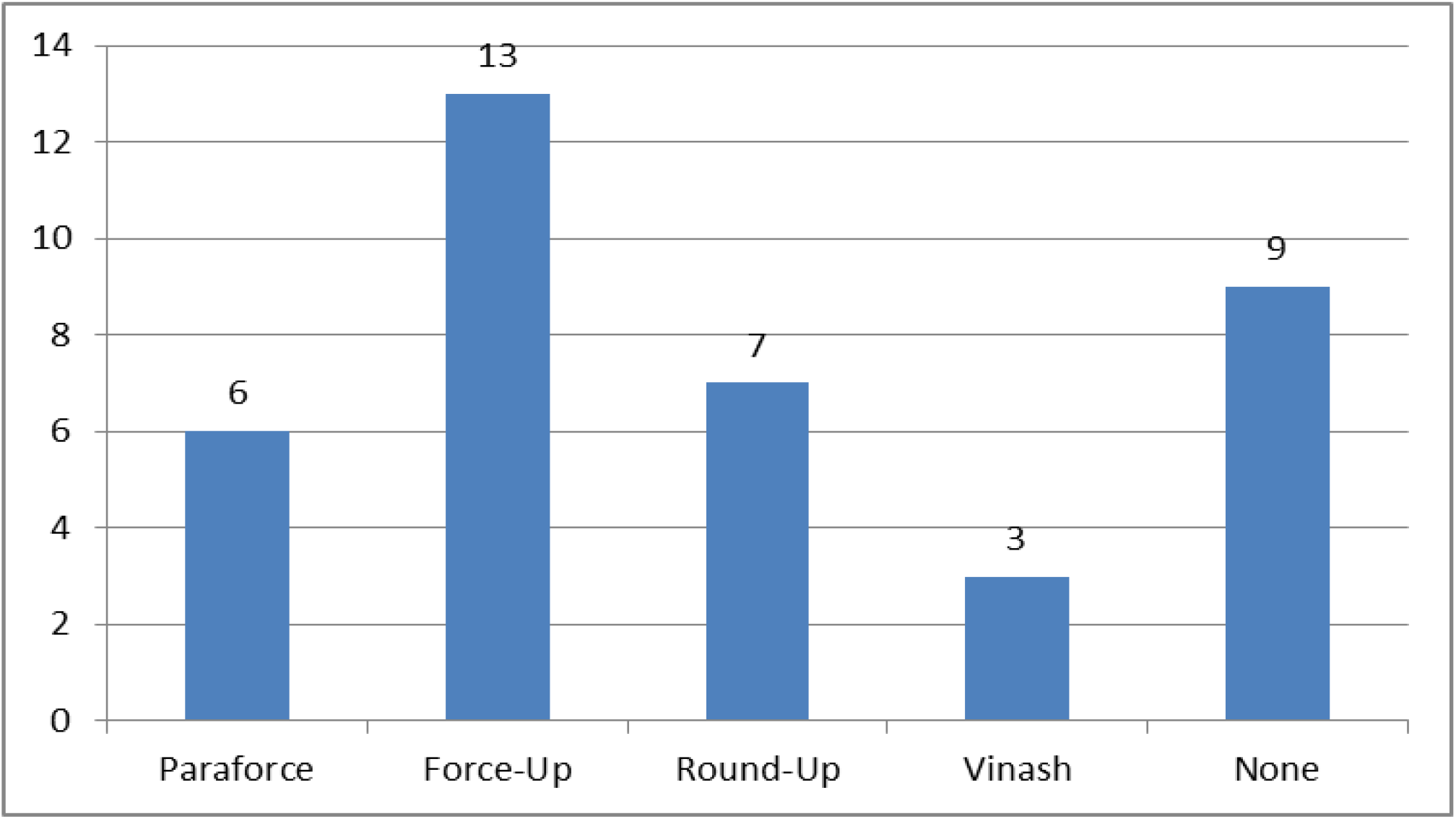
showing Distribution of Glyphosate-based Herbicides use in Fish Farms.

### Histopathological Lesions

There were mortalities at higher treatment concentrations with two (2) mortalities in the second treatment (T_2_) and five (5) mortalities in the third treatment (T_3_). The observable histopathological lesions mostly occurred at higher dose treatments (T_2_ and T_3_) while most of the organs in the lowest dose treatment (T_1_) appeared not affected. Histopathological lesions observed showed that livers and kidneys were mostly affected by glyphosate toxicity compared to the gills. Lesions observed in the gills include diffusely stunted and eroded secondary lamellae (Fig. 3) and severe congestion of the blood channel at the core of the primary lamellae (Fig. 4).

**Fig. 3:**
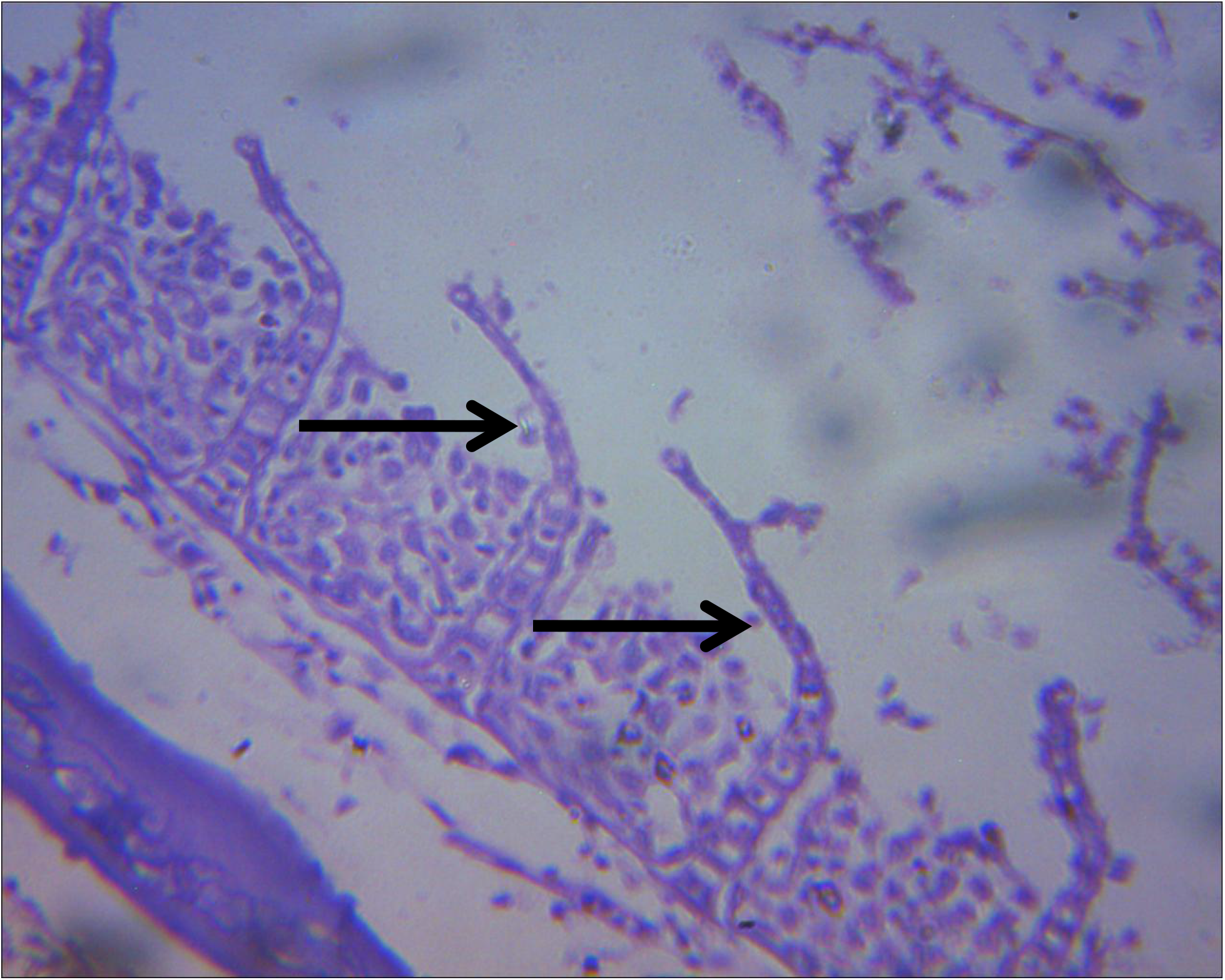
Diffusely Stunted and Eroded Secondary Lamellae of the Gills (Arrows) at First treatment concentration (0.15mls)

**Fig. 4:**
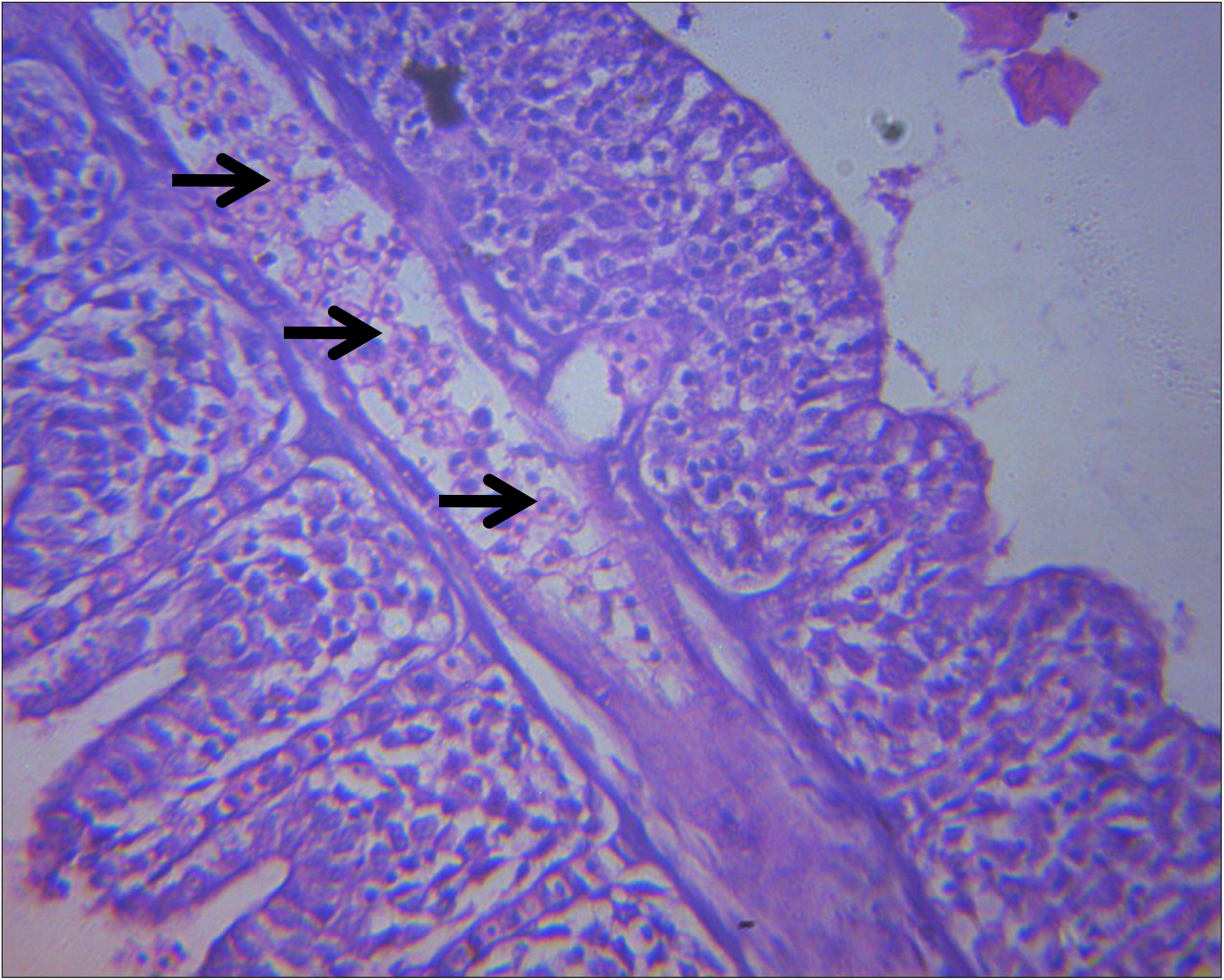
Severe Congestion of the Blood Channel at the Core of the Primary Lamellae (arrows) at second treatment concentration (0.225mls)

In the Liver samples, lesions observed include severe diffuse vacuolation of the hepatocytes at the first treatment concentration (Fig. 5), moderate to severe portal congestion, and mild diffuse vacuolation of hepatocytes at the second treatment concentration (Fig. 6). At the highest treatment dose of 0.3mls, there was moderate diffuse vacuolation of hepatocytes and severe portal congestion (Fig. 7).

**Fig. 5:**
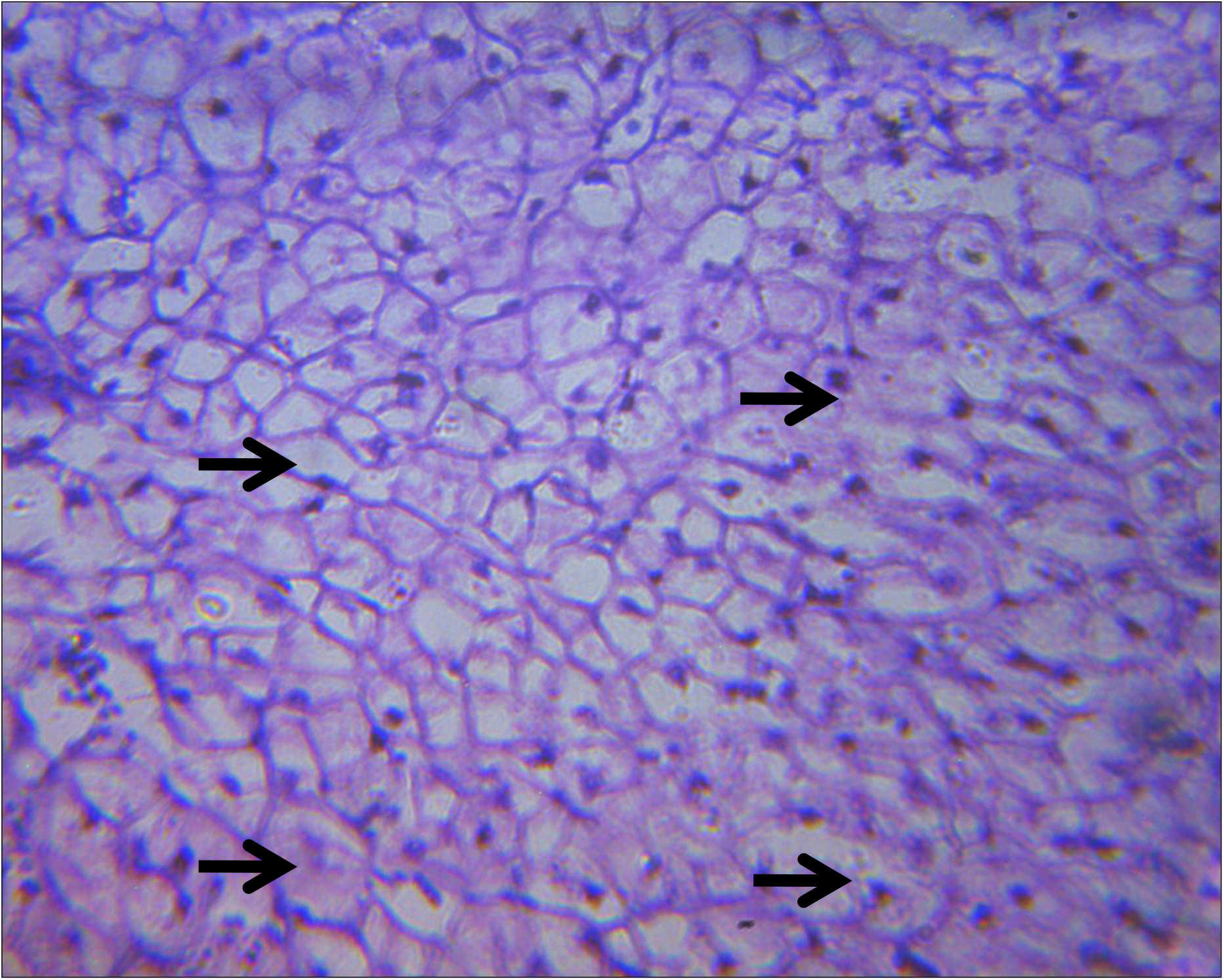
Severe Diffuse Vacuolation of the Hepatocytes (Arrows) at First Treatment Concentration (0.15mls)

**Fig. 6:**
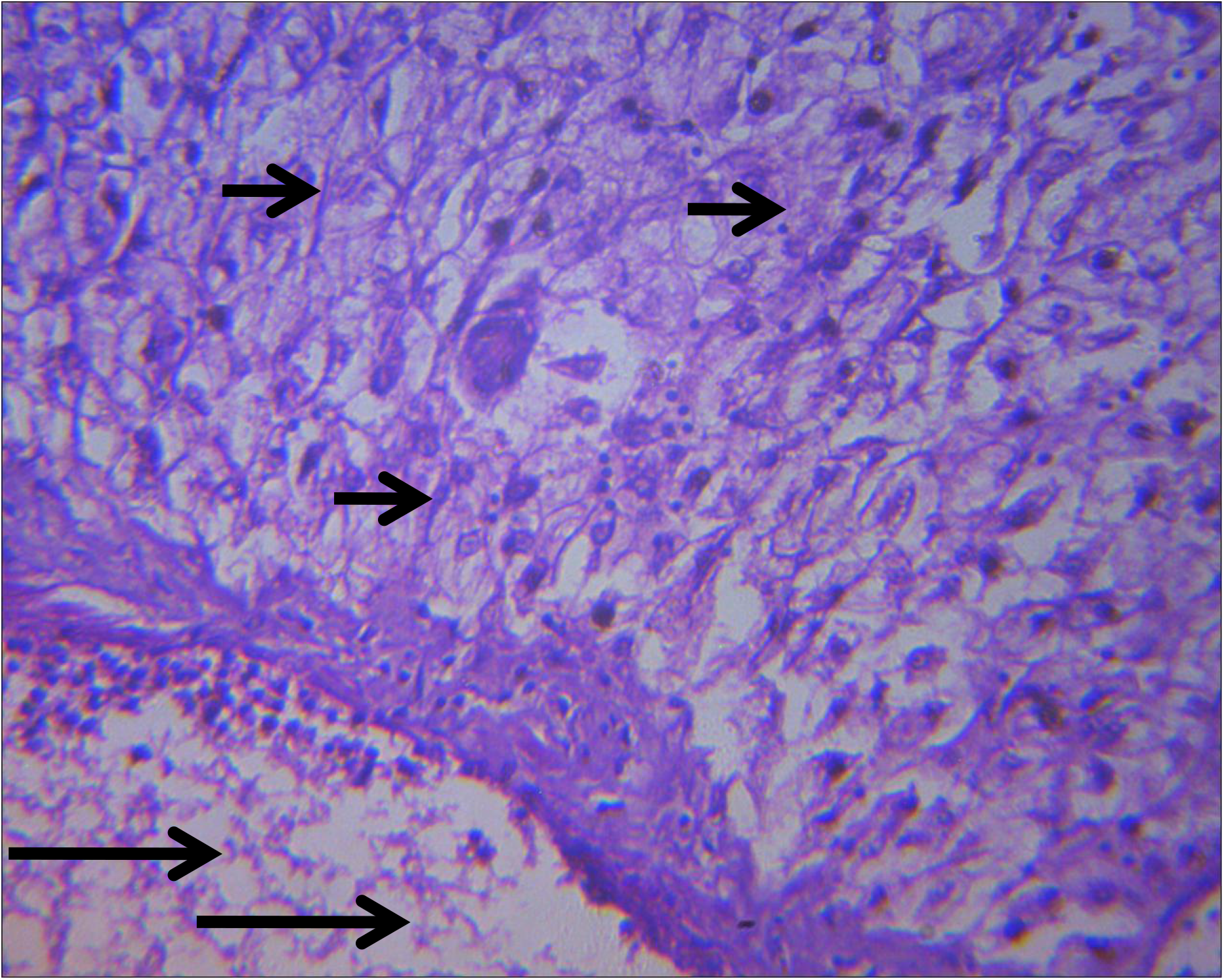
Moderate to Severe Portal Congestion (long arrows) and Mild Diffuse Vacuolation of Hepatocytes (short arrows) at Second Treatment Concentration (0.225mls)

**Fig. 7:**
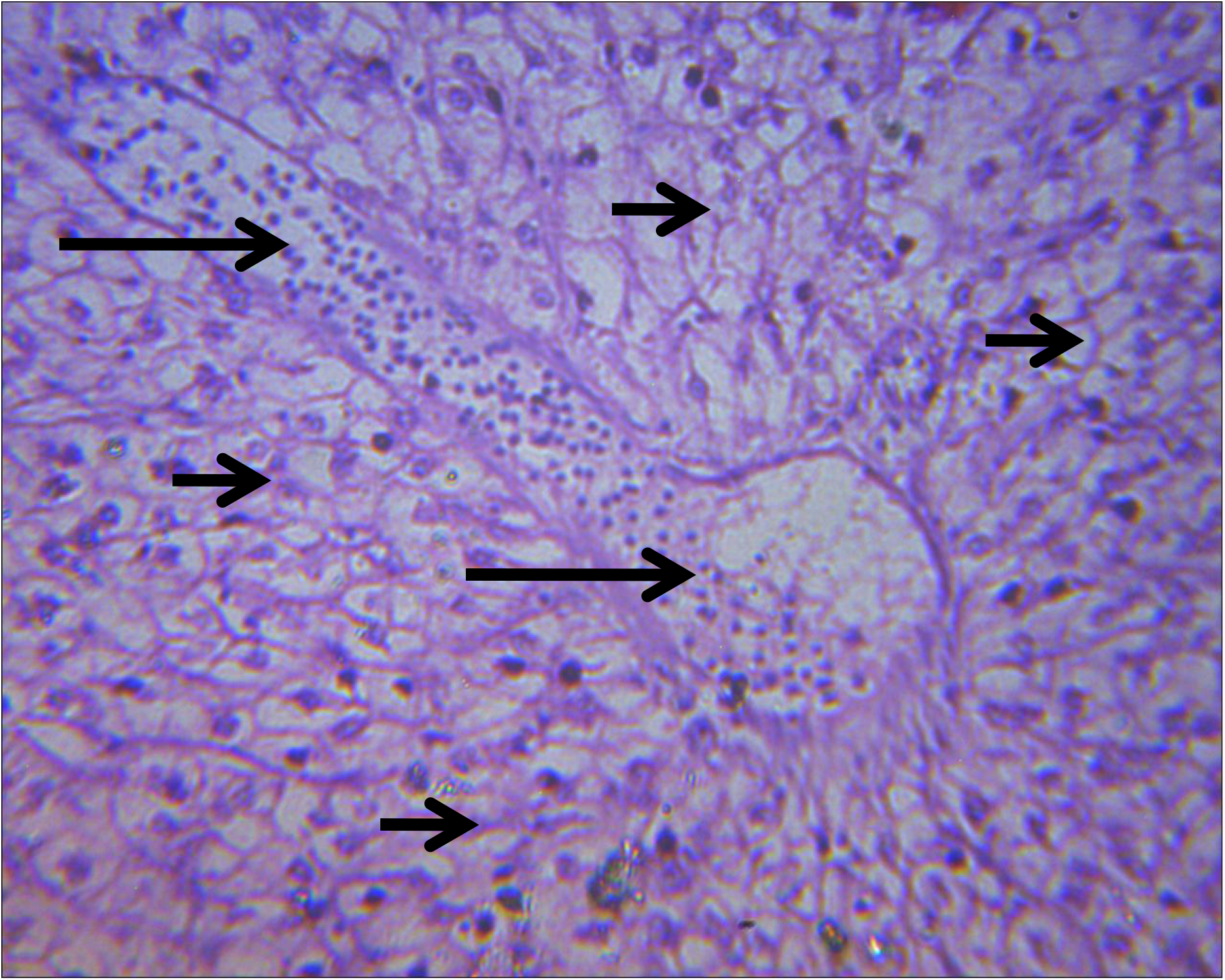
Moderate Diffuse Vacuolation of Hepatocytes and Severe Portal Congestion at Third Treatment Concentration (0.3mls)

The histopathological lesions observed in the kidney include mild to moderate congestion of the interstitium at second treatment concentration (Fig. 8) and focus of interstitial oedema within the parenchyma at third treatment concentration (Fig. 9).

**Fig. 8:**
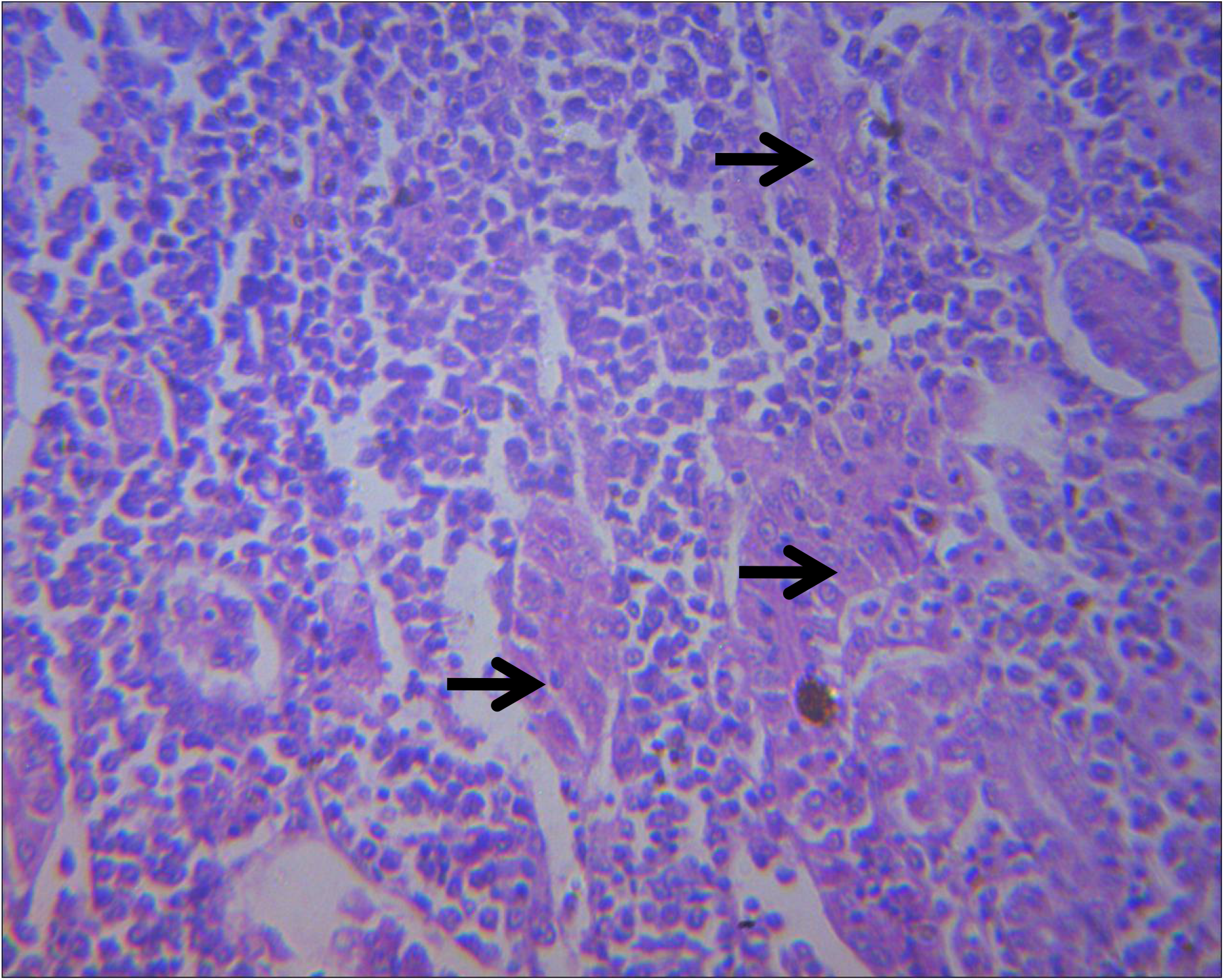
Mild to Moderate Congestion of the Interstitium at Second Treatment Concentration (0.225mls)

**Fig. 9:**
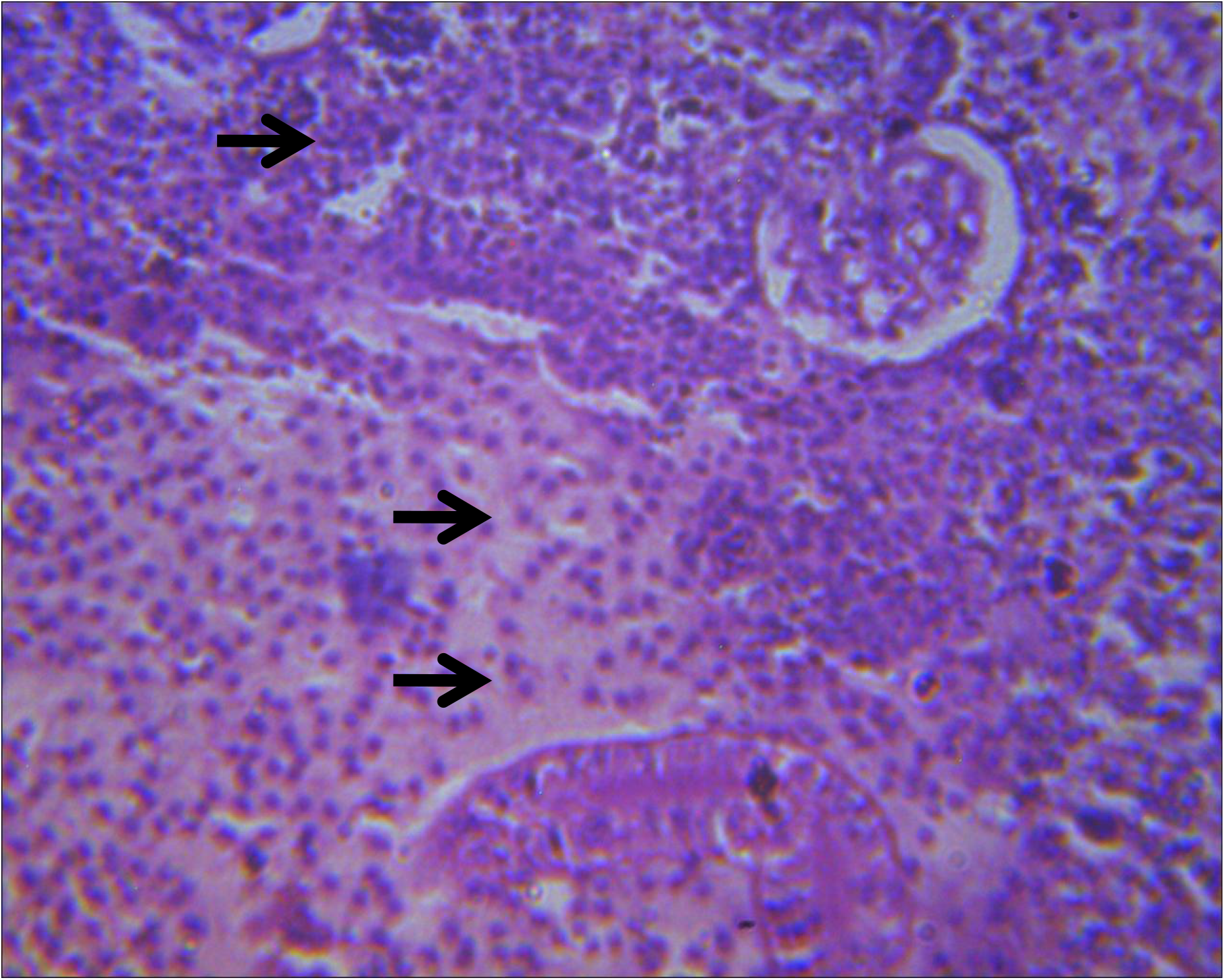
Focus of Interstitial Oedema within the Parenchyma at Third Treatment Concentration (0.3mls)

## MICRONUCLEUS ASSAY

Micronucleus assay result revealed normal fish nucleated red blood cells (Fig. 10) at both control and low treatment concentration (0.15mls) while the higher treatment (0.225mls and 0.3mls) concentrations showed micronucleus in the fish nucleated red blood cells (Figs. 11 and 12 respectively).

**Fig. 10:**
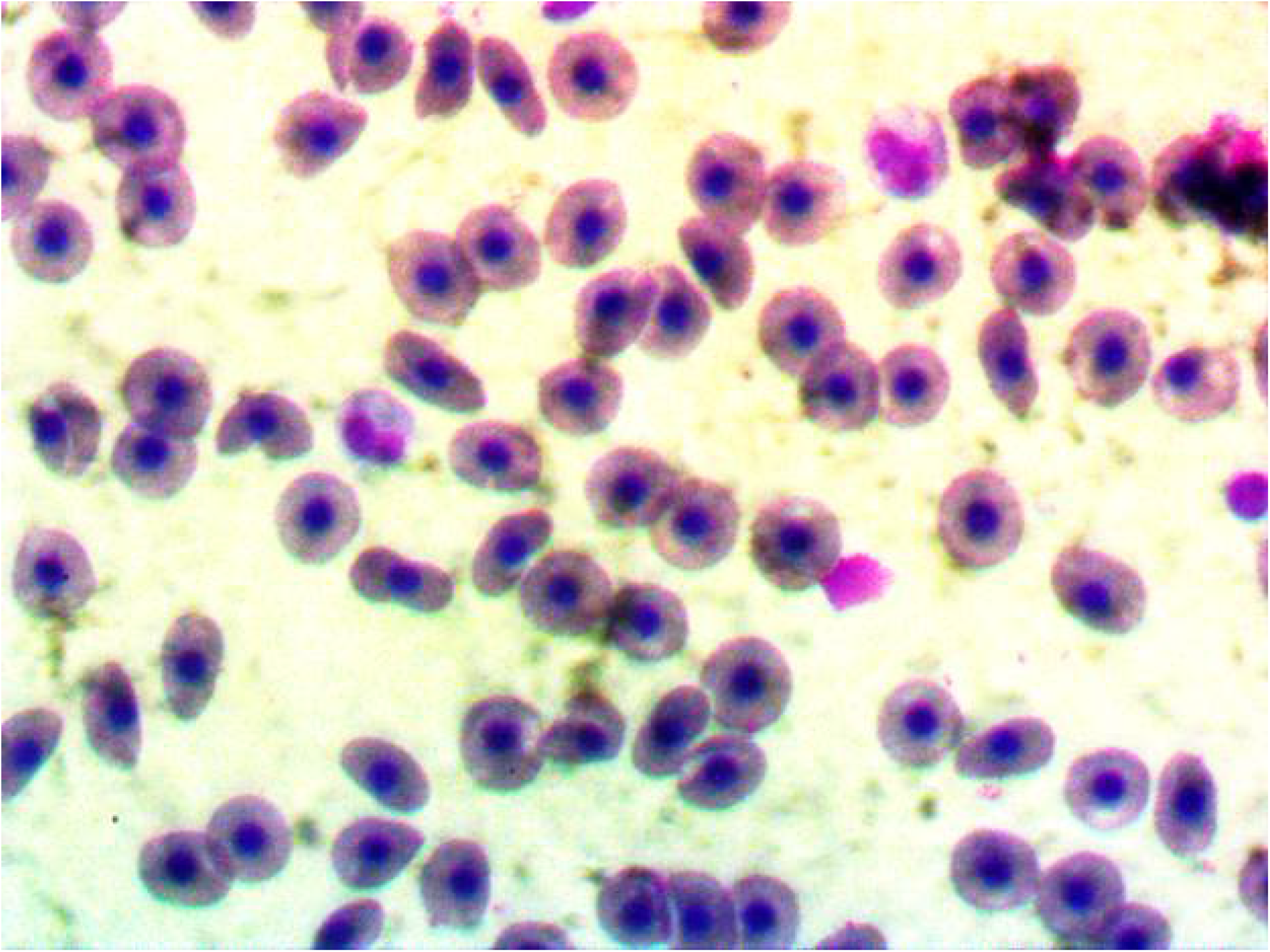
Photomicrograph showing Normal Fish Nucleated Red Blood Cell (Giemsa x1000)

**Fig. 11:**
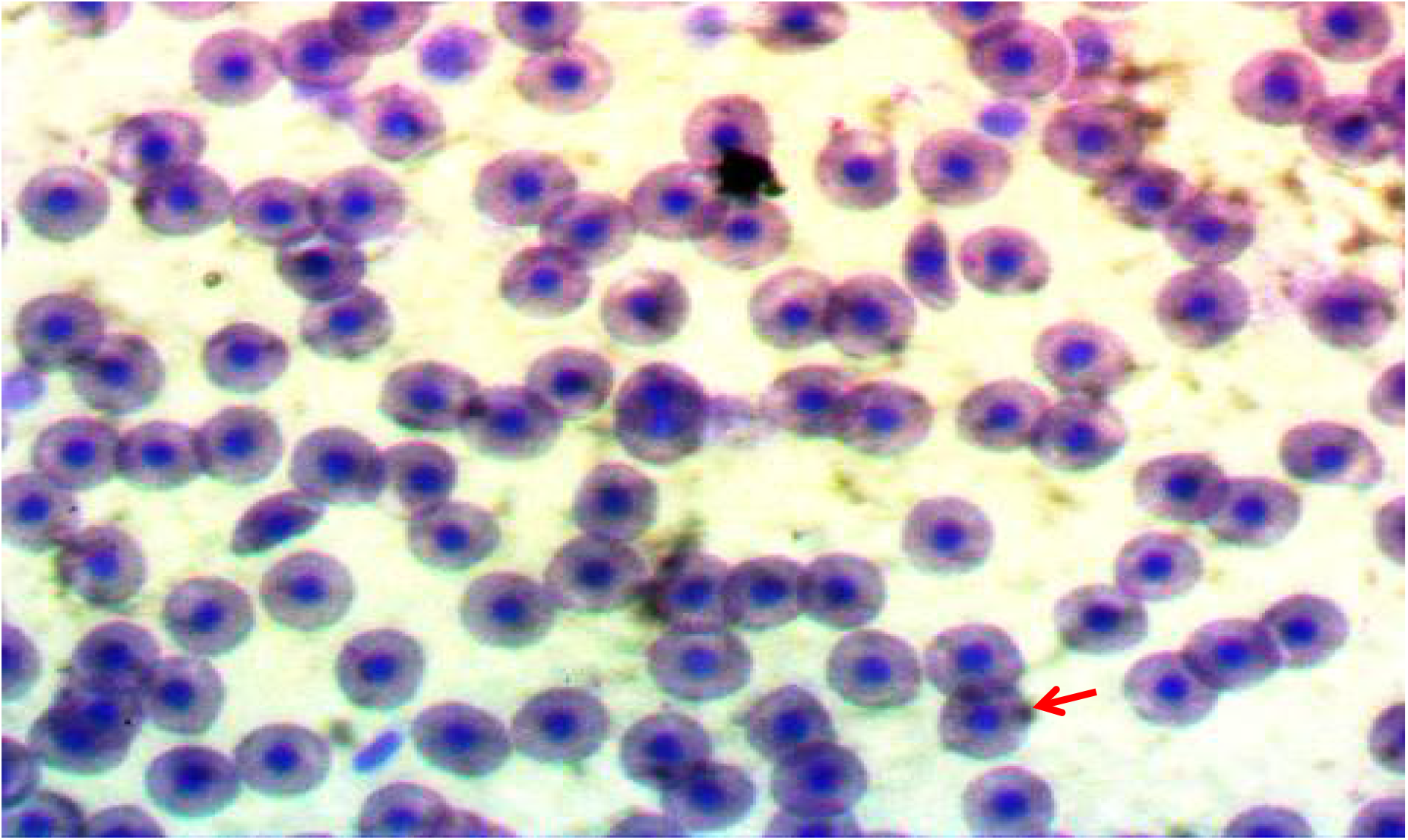
Photomicrograph showing Micronucleus (red arrow) in the Fish Nucleated Red Blood cell at 0.225ml Treatment Concentration (Giemsa x1000)

**Fig. 12:**
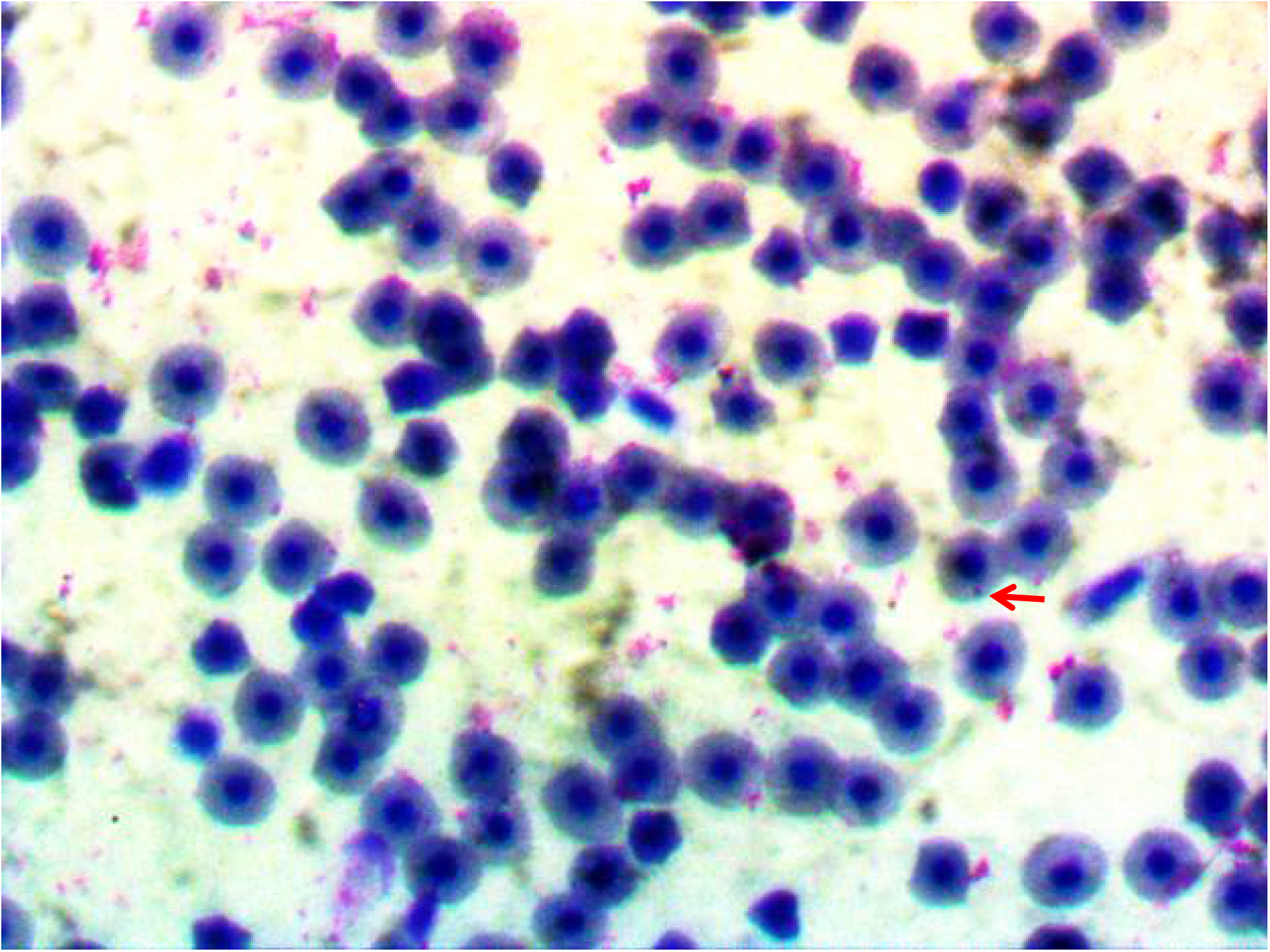
Photomicrograph showing Micronucleus (red arrow) in the Fish Nucleated Red Blood Cell at 0.3mls Treatment Concentration (Giemsa x1000)

## Discussion

Glyphosate is an herbicide that was introduced by Monsanto under the trade name Roundup in 1974 and in the last decade became the most widely used agricultural pesticide worldwide [21]. The above statement was validated by the responses of the respondents in this study where over two-thirds of the respondents (76.32%) were using one form of glyphosate or another in the control of weeds on the farm. Though some of the glyphosate products were originally used to control aquatic plants [21] but Fish farmers in Nigeria are using it in the control of weed around the pond and fish farms. Fish in the control experiment were calm and exhibited normal behavioural activities throughout the experimental periods while fish in the treatment groups showed erratic swimming, splashing, restlessness, movement from surface to bottom, sudden movement, and resting at the bottom of the container. The observed behavioural signs were similar to the several reports by Aguigwo, [22], Omoniyi *et al*., [23], Rahman *et al*., [24], Okayi *et al*., [25] and Sani and Idris, [26].

Histopathological lesions observed in collected tissues from the fish were predominantly pronounced in higher dose treatments which are similar to the report of Ayoola, [27] where he reported pronounced lesions at 96 hours exposure with accompanying mortality with an increase in the concentration of the glyphosate. The accompanied mortality pattern observed in this study is also in agreement with the reports of Ayoola, [18] and Sani and Idris, [26]. The observed high mortalities at the highest concentration of the herbicides demonstrated the observation of Fryer, [27] which stated that when a threshold of toxicants is reached, there is no drastic survival of the animal. When below the threshold, an animal will be in a tolerance zone while above the tolerance zone is the zone of resistance.

Observable histopathological lesions showed that livers and kidneys were mostly affected by the glyphosate toxicity compared to the gills which are in agreement with the report of Ayoola, [18]. The diffuse, stunted, and eroded secondary lamellae of the gills coupled with severe congestion of the blood channel at the core of the primary lamellae reported in this study were in line with previous findings of Nowak, [28], Neskovic *et al*., [29] and Ayoola, [18]. The presence of severe diffuse vacuolation of the hepatocytes, moderate to severe portal congestion, and mild diffuse vacuolation of hepatocytes in the liver samples were similar to the report of Ayoola, [18] where he reported that the most frequent types of degenerative changes encountered are hydropic degeneration, cloudy swelling, vacuolization, and focal necrosis. Risbourg and Bastide, [30] also reported an increase in the size of lipid droplets and vacuolization in the liver of fish exposed to atrazine herbicide. At the highest treatment dose of 0.3mls, there was moderate diffuse vacuolation of hepatocytes and severe portal congestion. In the present study, the kidney of *Clarias gariepinus* exposed to Force-Up showed mild to moderate congestion of the interstitium and focus of interstitial oedema within the parenchyma. These lesions were similar to the reports of Oulmi *et al*., [31], Omoniyi *et al*., [23], Rahman *et al*., [24].

Fish and aquatic invertebrates have been considered to be an excellent model for studying the toxic, mutagenic and carcinogenic potential of the water pollutants [32,33] as they play different roles in the trophic web such as undergoing bio-accumulation of environmental pollutants, biotransformation of xenobiotics through cytochrome P450-dependent oxidative metabolism and respond to mutagens at low concentration. The presence of micronucleus formation in the fish nucleated red blood cells at higher treatment (0.225mls and 0.3mls) concentrations in this study is also in support of a previous study carried out by Nwani et al., [34] which reported micronucleus formation in erythrocytes of *C. punctatus* after exposure to carbosulfan (0.07, 0.13 and 0.20 mg L^1^), glyphosate (8.1, 16.3 and 24.4 mg L^1^) and atrazine (10.6, 21.2 and 31.8 mg L^1^) for 96 hours.

## Conclusion

Glyphosate is used extensively in aquaculture, and it’s the world’s most used herbicide. The results and the available information on glyphosate toxicity and its formulations on different groups of cultured catfish in this study revealed that they are menacing and unsafe to the aquatic environment, and cause histopathological changes in different organs especially, at higher concentrations and particularly, in livers, kidneys, and gills.

Despite the benefits derived from the use of glyphosate-based herbicides, the result of this work, like many others, portends that herbicides can potentially harm the aquatic environment, human health and can even alter the world food chain cycle. Glyphosate residue could reach humans and animals through feed consumption and contact with or use of contaminated water. Unknown impacts of glyphosate on human and animal health warrant further investigations of glyphosate residues in vertebrates and other non-target organisms). Finally, the study has therefore established that glyphosate-based herbicides are toxic to *Clarias gariepinus* (African Catfish) at high concentrations.

Farmers should also be discouraged not to use high concentrations of herbicides around fish ponds. Most of the world’s population depends on fish as another major alternative to a protein source, then the agricultural and aquaculture communities should therefore be conscious of the potential adverse effects of this pesticide and several others. To protect the water quality and safety of aquatic animals, Government should therefore enact policies that will guide against the use of dangerous herbicides in the control of terrestrial and aquatic weeds. Also, farming activities should not be too close to water bodies especially, farms that rely on chemical control of weeds. This is to prevent the water body from the residue of herbicides that would have washed down to the water.

## Author Contributions

**Conceptualization:** Olanike K. Adeyemo, Selim A. Alarape

**Sample Collection:** Emmanuel Oyindamola Adebiyi

**Methodology:** Selim A. Alarape, Emmanuel Oyindamola Adebiyi

**Formal analysis:** Selim A. Alarape

**Supervision**: Olanike K. Adeyemo, Selim A. Alarape

**Validation:** Olanike K. Adeyemo.

**Writing:– original draft:** Selim A. Alarape

**Writing – review & editing:** Olanike K. Adeyemo, Selim A. Alarape.

